# Neuromodulated Equilibrium Learning

**DOI:** 10.64898/2026.08.28.747901

**Authors:** Yoshimasa Kubo

**Affiliations:** Department of Computer Science, Lakehead University, Thunder Bay, ON, Canada

## Abstract

Equilibrium Propagation (EP) is a biologically motivated alternative to backpropagation but requires free and nudged equilibrium phases, raising questions about the biological plausibility of phase-separated learning. We introduce Neuromodulated Equilibrium Learning (NEL), a single-equilibrium reward-based framework inspired by Attention-Gated Brain Propagation and neuromodulatory signaling. After reaching a free equilibrium, NEL propagates a reward-dependent modulatory signal derived from the selected action through reciprocal feedback connections to drive local synaptic updates, eliminating both the reward-nudged equilibrium and contrastive update. We evaluate NEL on MNIST, Fashion-MNIST, and CIFAR-10 using multilayer perceptrons and convolutional networks, with symmetric three-phase REP and standard EP as multi-phase baselines. NEL approaches three-phase REP in shallow architectures, with gaps of 0.79 and 1.31 percentage points on MNIST and one-hidden-layer Fashion-MNIST, while larger gaps of 3.13 and 6.21 percentage points emerge in the deeper Fashion-MNIST and CIFAR-10 settings. A direct cost feedback control using the full supervised output error also remains below three-phase EP, indicating that sparse reward information alone does not explain the performance gap between single-equilibrium and multi-phase learning. These results show that effective reward-driven learning can arise from a single free equilibrium while suggesting that computations introduced by additional nudged relaxations become increasingly important in more challenging architectures.

## 1 Introduction

Equilibrium Propagation (EP) (Scellier & Bengio, 2017, 2019; Ernoult et al., 2019; Laborieux et al., 2021; Laborieux & Zenke, 2022) is a biologically motivated alternative to backpropagation (BP) (Rumelhart et al., 1986) that enables synaptic updates from quantities locally available at the synapse. Recent advances have further shown that improved variants of EP can train convolutional neural networks with performance approaching that of BP (Laborieux et al., 2021).

A defining feature of EP is its use of distinct dynamical phases. During the free phase, the network relaxes without an explicit teaching signal, whereas during subsequent nudged relaxation, a weak task-dependent perturbation drives the network toward a nearby equilibrium. Synaptic updates are obtained from differences in neural activities across these equilibria. Because the resulting learning rule can be expressed using locally available pre- and postsynaptic activities, EP provides an appealing framework for biologically plausible credit assignment (Rumelhart et al., 1986; Sutton et al., 1998; Roelfsema & Ooyen, 2005; Friedrich et al., 2011; Bengio, 2014; Lee et al., 2014; Roelfsema & Holtmaat, 2018). However, phase-separated learning may require neural activity to be retained across temporal periods or plasticity to be coordinated according to the current phase.

Several biologically motivated approaches have attempted to reduce these temporal requirements. Dendritic cortical microcircuits can support supervised error-driven learning without explicitly separated phases (Sacramento et al., 2017), Continual Equilibrium Propagation (C-EP) updates synaptic weights continuously during the nudged phase (Ernoult et al., 2020), and Latent Equilibrium (LE) uses prospective neuronal states to avoid waiting for complete relaxation (Haider et al., 2021). These approaches reduce phase separation or relaxation requirements in different ways, but it remains unclear whether equilibrium-based learning driven by sparse reward information can eliminate the reward-nudged equilibrium itself.

Reward-based learning provides a complementary approach to biologically plausible credit assignment. Attention-Gated Brain Propagation (BrainProp) (Pozzi et al., 2020) introduced a neuromodulatory mechanism in which action-dependent rewards influence synaptic plasticity through feedback pathways. Reward-based Equilibrium Propagation (REP) (Kubo, 2026b) extended this principle to recurrent equilibrium networks, enabling learning from selected actions and scalar rewards rather than a complete supervised output error. REP, however, retains reward-nudged relaxation and contrastive updates inherited from EP.

Here, we introduce Neuromodulated Equilibrium Learning (NEL), a single-equilibrium framework for reward-driven learning. After the network relaxes to a free equilibrium, an action is selected and evaluated by a scalar reward. The resulting reward-dependent modulatory signal is propagated through reciprocal feedback connections and combined with locally available neuronal activities to drive synaptic updates. NEL therefore eliminates reward-nudged relaxation and contrastive comparison between equilibrium states while also avoiding activation derivatives in its modulatory pathway.

We evaluate NEL on MNIST (LeCun & Cortes, 2005), Fashion-MNIST (Xiao et al., 2017), and CIFAR-10 (Krizhevsky, 2009) using multilayer perceptrons and convolutional neural networks. Symmetric three-phase REP and standard three-phase EP provide multi-phase baselines, while truncated BPTT following Laborieux et al. (2021) provides a gradient-based reference that propagates supervised credit through the free-relaxation trajectory without additional nudged equilibrium phases. NEL approaches three-phase REP in relatively shallow architectures, whereas larger gaps emerge in deeper and convolutional networks. We additionally introduce a Direct Cost Feedback (DCF) control that replaces NEL’s sparse reward signal with the full supervised output error while retaining the same free-equilibrium dynamics and local feedback-based learning structure. DCF does not systematically improve over NEL and remains below standard three-phase EP across the evaluated settings, indicating that sparse reward information alone is insufficient to explain the performance gap. In contrast, BPTT recovers much of the performance of three-phase learning without additional nudged equilibria. Together, these results demonstrate that useful reward-driven learning can arise from a single free equilibrium while suggesting that effective propagation of teaching information through the network dynamics, rather than the number of equilibrium phases alone, is a key determinant of performance.

## 2 Methods

### 2.1 Equilibrium Propagation

Equilibrium Propagation (EP) is a learning framework for energy-based recurrent neural networks in which neuronal states evolve toward equilibrium through recurrent dynamics (Scellier & Bengio, 2017, 2019). Let **x** denote the input, **s** the collection of neuronal states, **t** the target, and *θ* the trainable synaptic parameters. The network is characterized by an energy function *E*(**s**, *θ*, **x**). When a task-dependent cost *C*(**s, t**) is introduced, the total energy *F* is defined as

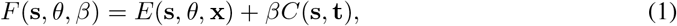

where *β* controls the strength of the teaching perturbation. The neuronal states evolve according to the continuous-time dynamics

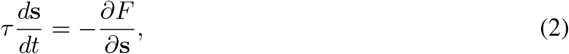

where *τ* denotes a neuronal time constant.

In the standard two-phase formulation of EP, learning consists of a *free phase* followed by a *weakly clamped phase*, also referred to as the nudged phase. During the free phase, the input **x** is fixed and no teaching signal is applied. Setting *β* = 0 in Eq. 2 gives

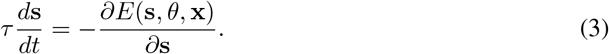

The network evolves until its neuronal states approach a free equilibrium, denoted by **s**^0^, which represents the network response before teaching information is introduced.

Starting from **s**^0^, the weakly clamped phase introduces a small nonzero value of *β*. The cost weakly perturbs the network, and the dynamics become

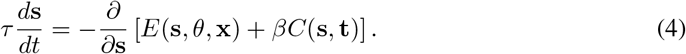

The network relaxes toward a nearby weakly clamped equilibrium, denoted by **s**^*β*^. The synaptic update is obtained from the contrast between the two equilibrium configurations,

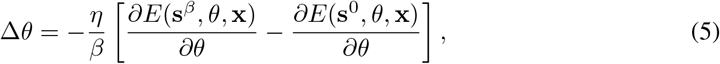

where *η* denotes the learning rate. Thus, EP learning is based on the change in the network equilibrium induced by a weak teaching perturbation.

### 2.2 Attention-Gated Brain Propagation

Attention-Gated Brain Propagation (BrainProp) (Pozzi et al., 2020) provides a reward-based approach to credit assignment in which a network learns from the outcome of its selected action rather than from a complete supervised target vector. For a given input, the network produces an output activity vector **y**, where each output unit represents a possible class or action.

BrainProp uses a Max–Boltzmann action-selection mechanism. With probability 1 − *ϵ*, the action associated with the most active output unit is selected,

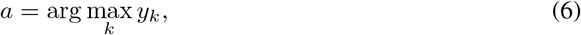

whereas with probability *ϵ*, an exploratory action is sampled according to

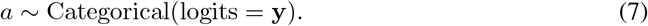

After selecting an action, the network receives a binary reward,

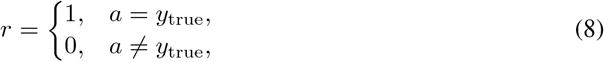

where *y*_true_ denotes the correct class. When an incorrect action is selected, the correct output class is not directly revealed to the network.

The activity *y*_*a*_ of the selected output unit is treated as the predicted reward associated with action *a*, giving the selected-output objective

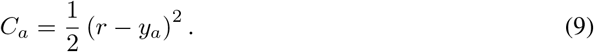

The corresponding reward-dependent output signal is

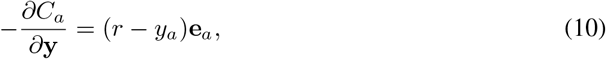

where **e**_*a*_ is the one-hot basis vector associated with the selected action. Thus, the direct reward-dependent teaching signal is nonzero only for the selected output unit.

In BrainProp, feedback connections propagate action-specific credit information toward earlier layers, where it modulates synaptic plasticity. This reward-based credit-assignment principle forms the basis for its subsequent integration with equilibrium-based learning.

### 2.3 Reward-Based Equilibrium Propagation

Reward-Based Equilibrium Propagation (REP) (Kubo, 2026b) combines the reward-based learning principle of BrainProp with the recurrent equilibrium dynamics of EP. REP adopts the action-selection and selected-output reward formulation in Eqs. 6–10, while using equilibrium dynamics to propagate the resulting reward-dependent perturbation through the network.

REP first performs the same free phase as conventional EP, evolving according to Eq. 3 until reaching the free equilibrium **s**^0^. Let **y**^0^ denote the output activity at this equilibrium. An action *a* is selected from **y**^0^ using the Max–Boltzmann mechanism, and the corresponding binary reward *r* is obtained. The selected action and reward are determined once from the free equilibrium and remain fixed throughout the subsequent reward-nudged relaxations.

REP defines a reward-dependent cost only for the selected output unit,

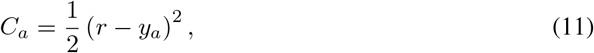

where *y*_*a*_ denotes the current activity of the selected output unit. Introducing a nonzero nudging factor *β* modifies the neuronal dynamics to

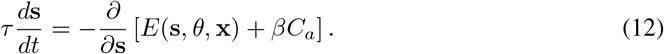

Because *C*_*a*_ depends only on the selected output unit, the direct reward-dependent teaching signal is

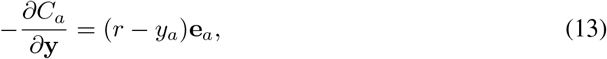

where **e**_*a*_ is the one-hot basis vector corresponding to the selected action. At the free equilibrium, this signal is 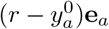. Although the perturbation is applied directly only to the selected output unit, recurrent interactions propagate its influence toward hidden layers during the nudged dynamics.

In this study, all REP experiments use the symmetric three-phase formulation of EP (Laborieux et al., 2021). Starting from the free equilibrium **s**^0^, the network is relaxed under a positive reward-dependent perturbation +*β* and a negative reward-dependent perturbation −*β*, producing two reward-nudged equilibria, **s**^+*β*^ and **s**^−*β*^. The same action *a* and binary reward *r* obtained from the free equilibrium are used in both nudged phases.

The centered parameter update is

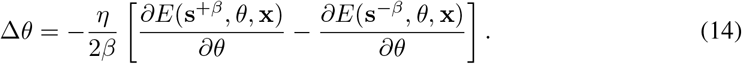

Thus, REP combines the action-selection and selected-output reward mechanism of BrainProp with the recurrent credit propagation and contrastive learning mechanism of EP. The reward signal is sparse at the output because it directly constrains only the selected action, but the positive and negative reward-nudged relaxations transform this signal through the recurrent network before the synaptic update is computed.

### 2.4 Neuromodulated Equilibrium Learning

Building on the reward-dependent learning principle of BrainProp and its integration with equilibrium dynamics in REP, we propose *Neuromodulated Equilibrium Learning* (NEL). NEL retains the free-equilibrium computation but eliminates the subsequent reward-nudged relaxation and contrastive comparison between equilibrium states.

For each input, NEL performs the free relaxation defined in Eq. 3 until reaching the free equilibrium **s**^0^. An action and its corresponding reward are then obtained using the BrainProp-inspired mechanism in Eqs. 6–8. Whereas REP injects the reward-dependent signal into the neuronal dynamics and relaxes toward an additional reward-nudged equilibrium, NEL uses this signal directly as a neuromodulatory factor for synaptic plasticity.

#### 2.4.1 Neuromodulatory Feedback

Let **y**^0^ denote the output activity at the free equilibrium. For the selected action *a*, the output-layer modulatory signal is

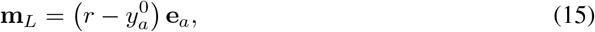

corresponding to the selected-output reward signal evaluated at the free equilibrium.

Rather than injecting this signal into the neuronal dynamics, NEL propagates it directly toward earlier layers through reciprocal feedback connections. Let *W*_*l*_ denote the synaptic weight matrix connecting layer *l* − 1 to layer *l*. The modulatory signal is propagated recursively according to

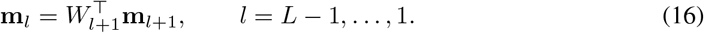

Unlike the error gradient propagated by conventional backpropagation, this modulatory pathway does not include derivatives of the neuronal activation functions.

#### 2.4.2 Local Synaptic Plasticity

The propagated modulatory signal is combined with neuronal activities at the free equilibrium to determine synaptic plasticity. Let 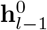 and 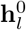 denote the presynaptic and postsynaptic equilibrium activities associated with *W*_*l*_, respectively, with 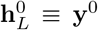 for the output layer. For a mini-batch containing *B* samples, the synaptic weight update is

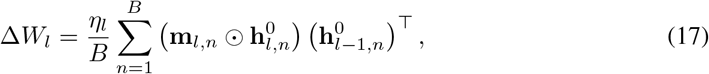

where ⊙ denotes element-wise multiplication and *η*_*l*_ denotes the layer-specific learning rate. The corresponding bias update is

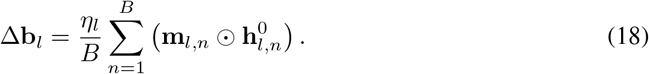

The resulting rule has the form of a three-factor plasticity rule, in which presynaptic and postsynaptic activities are combined with a reward-dependent modulatory signal (Frémaux & Gerstner, 2016; Roelfsema & Holtmaat, 2018). The local update therefore follows the reward-modulated credit-assignment principle underlying BrainProp while using neuronal activities obtained from the free equilibrium. Importantly, all layer-wise updates are computed from the same free-equilibrium state and pre-update synaptic parameters before any parameter is modified.

NEL thus replaces REP’s reward-nudged relaxation and contrastive update with direct neuromodulatory feedback and local synaptic plasticity. Unlike REP, where the teaching perturbation is transformed by recurrent dynamics during the nudged phase, NEL propagates the reward-dependent signal directly through the reciprocal feedback pathway. NEL is not assumed to be mathematically equivalent to the nudged dynamics of EP; their relationship is discussed in Appendix A.

#### 2.4.3 Convolutional Extension

The same principle extends to convolutional networks through reciprocal convolutional operations. During free-equilibrium relaxation, bottom-up interactions use convolution followed by max pooling, while the reciprocal top-down pathway uses the stored pooling switches for max unpooling followed by transpose convolution with tied forward kernels. The reward-dependent modulatory signal is propagated through the same reciprocal pathway. At each convolutional layer, it is combined element-wise with the postsynaptic equilibrium activity, unpooled using the stored pooling switches, and correlated with the corresponding presynaptic activity to determine the local weight update. As in the fully connected formulation, convolutional NEL requires neither activation derivatives nor an additional reward-nudged equilibrium.

### 2.5 Model and Dataset Specifications

We evaluated NEL, REP, standard Equilibrium Propagation (EP), and backpropagation through time (BPTT) on MNIST, Fashion-MNIST (FMNIST), and CIFAR-10 using multilayer perceptrons (MLPs) and convolutional neural networks (CNNs). Direct Cost Feedback (DCF) was additionally evaluated as a control. DCF retained the same free-equilibrium dynamics, reciprocal feedback pathway, and local plasticity structure as NEL, while replacing the sparse reward-based teaching signal with the full supervised output signal **t** − **y**^0^.

MNIST and FMNIST images were flattened into 784-dimensional vectors for the MLP experiments. CIFAR-10 was evaluated using a two-layer convolutional network without an intermediate fully connected hidden layer. The same dataset preprocessing was used across methods, and no data augmentation was applied.

Unless otherwise stated, all output layers used the hard-sigmoid activation, the mini-batch size was 64, and results were averaged over six random seeds. NEL and DCF used only free-equilibrium relaxation. REP and EP used the symmetric three-phase formulation, with positive and negative nudged relaxations around the free equilibrium. REP used reward-based nudging of the selected output, whereas standard EP used the full supervised target during the nudged phases. Following Laborieux et al. (2021), BPTT used the same free relaxation dynamics and propagated supervised gradients through the final 12 relaxation steps using truncated backpropagation through time.

Detailed architectures, learning rates, relaxation lengths, activation functions and bounds, dataset preprocessing, and training durations are provided in Appendix B.

Code for reproducing the experiments will be made publicly available.

## 3 Results

### 3.1 Classification Performance

Table 1 summarizes the maximum test accuracy of NEL, Direct Cost Feedback (DCF), BPTT, three-phase REP, and three-phase EP across the evaluated architectures. We primarily compare NEL with REP as reward-based learning rules, while the DCF–EP comparison provides a complementary supervised control. BPTT provides an additional gradient-based reference in which supervised gradients are propagated through the final 12 steps of the free relaxation dynamics. REP and EP use symmetric three-phase relaxation, whereas NEL, DCF, and BPTT do not require additional nudged equilibrium phases.

**Table 1:** Maximum test accuracy (%, mean ± std over six random seeds) for NEL, Direct Cost Feedback (DCF), BPTT, three-phase REP, and three-phase EP.

| Dataset | Architecture | NEL | DCF | BPTT | REP (3-phase) | EP (3-phase) |
| --- | --- | --- | --- | --- | --- | --- |
| MNIST | 784-512-10 | 97.62 ± 0.05 | 97.65 ± 0.03 | 98.19 ± 0.04 | 98.41 ± 0.03 | 98.24 ± 0.04 |
| FMNIST | 784-1024-10 | 88.42 ± 0.13 | 88.65 ± 0.15 | 90.15 ± 0.13 | 89.73 ± 0.11 | 90.04 ± 0.11 |
| FMNIST | 784-512-256-10 | 86.36 ± 0.29 | 87.83 ± 0.36 | 89.63 ± 0.11 | 89.49 ± 0.10 | 89.29 ± 0.11 |
| CIFAR-10 | Conv64-Conv128-10 | 65.07 ± 0.51 | 64.58 ± 0.22 | 70.57 ± 0.42 | 71.28 ± 0.22 | 72.46 ± 0.44 |

On MNIST, NEL achieved 97.62 ± 0.05%, compared with 98.41 ± 0.03% for three-phase REP, corresponding to a gap of 0.79 percentage points. On the one-hidden-layer FMNIST network, the corresponding gap was 1.31 percentage points. Thus, NEL remained relatively close to three-phase REP in both shallow MLP settings despite learning from only a single free-equilibrium relaxation.

Larger differences emerged with increasing architectural complexity. The NEL–REP gap increased to 3.13 percentage points on the two-hidden-layer FMNIST network and to 6.21 percentage points on CIFAR-10. A similar comparison between DCF and standard three-phase EP showed gaps of 0.59, 1.39, 1.46, and 7.88 percentage points on MNIST, one-hidden-layer FMNIST, two-hidden-layer FMNIST, and CIFAR-10, respectively.

BPTT substantially reduced these gaps while still operating without additional nudged equilibrium phases. It achieved 98.19 ± 0.04% on MNIST, only 0.05 percentage points below three-phase EP, and reached 90.15 ± 0.13% and 89.63 ± 0.11% on the one- and two-hidden-layer FMNIST networks, respectively, slightly exceeding three-phase EP in both cases. On CIFAR-10, BPTT achieved 70.57 ± 0.42%, remaining 1.89 percentage points below three-phase EP but substantially above both NEL and DCF.

Taken together, these results suggest that the performance advantage of the multi-phase methods cannot be attributed solely to the presence of additional nudged equilibrium relaxations. The strong BPTT results show that supervised credit propagated through the free relaxation trajectory can recover much of the performance of three-phase learning without introducing additional equilibrium phases. At the same time, the remaining gap on CIFAR-10 indicates that the effectiveness of this credit-assignment mechanism may depend on architectural complexity. These results therefore point more broadly to the importance of how teaching information is propagated through the network dynamics, rather than simply whether learning uses single- or multi-phase relaxation.

### 3.2 Direct Cost Feedback Control

To examine whether the performance gap between NEL and REP could be explained primarily by the limited information contained in NEL’s reward-based teaching signal, we evaluated Direct Cost Feedback (DCF) as a control. DCF retained the same free-equilibrium relaxation, reciprocal feedback pathway, and local plasticity structure as NEL, but replaced the sparse selected-action reward signal with the full supervised output signal **t** − **y**^0^, where **t** denotes the one-hot target vector. This also provides a complementary comparison with standard three-phase EP, which uses the same full supervised target but propagates it through additional nudged relaxations.

As shown in Table 1, DCF produced only small changes relative to NEL on MNIST, the one-hidden-layer FMNIST model, and CIFAR-10. A larger difference appeared for the two-hidden-layer FMNIST network, where DCF improved over NEL by 1.47 percentage points. However, DCF remained below three-phase EP across all evaluated settings, with gaps of 0.59, 1.39, 1.46, and 7.88 percentage points on MNIST, one-hidden-layer FMNIST, two-hidden-layer FMNIST, and CIFAR-10, respectively. In the two-hidden-layer FMNIST setting, DCF also remained 1.66 percentage points below three-phase REP. Stable DCF training in this setting required a learning rate of 0.08 per layer; the NEL-matched learning rate of 0.12 produced late-training instability, as detailed in Appendix B.

Taken together, these results show that replacing NEL’s sparse reward-based teaching signal with a full supervised output error does not systematically improve performance. Richer output supervision provided a clearer benefit in the deeper FMNIST MLP, but did not eliminate the performance gap and provided no advantage over NEL on CIFAR-10. Together with the strong BPTT results, this indicates that limited output information alone is insufficient to explain NEL’s performance gap. Rather, the effectiveness of the mechanism used to propagate teaching information through the network dynamics appears to be an important factor.

### 3.3 Generalization Behavior

We next compared the training dynamics of NEL and DCF using the epoch-wise generalization gap,

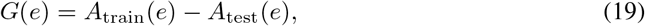

where *A*_train_(*e*) and *A*_test_(*e*) denote the training and test accuracies at epoch *e*, respectively. Figure 1 shows the mean gap across six random seeds, with shaded regions indicating variability across seeds.

**Figure 1:**
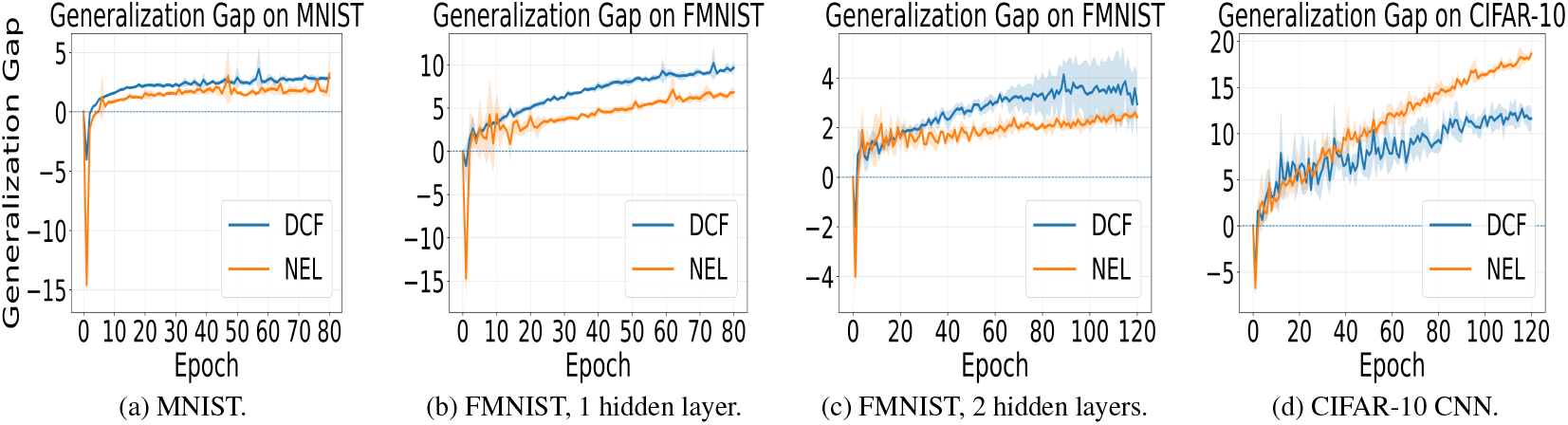
Generalization gap, defined as training accuracy minus test accuracy, for NEL and Direct Cost Feedback (DCF). Curves show the mean over six random seeds, with shaded regions indicating variability across seeds. NEL exhibited a smaller gap than DCF during later training across the evaluated MLP architectures, whereas this pattern did not extend to the CIFAR-10 CNN.

Across the three MLP experiments, NEL exhibited a smaller generalization gap than DCF during later training. This pattern was observed on MNIST and both FMNIST architectures, including the two-hidden-layer model for which DCF achieved higher maximum test accuracy than NEL. These results indicate that sparse reward-modulated learning and full supervised cost feedback produce different training behavior and are consistent with a possible regularization-like effect of NEL in the MLP settings considered here.

The pattern did not extend to the CIFAR-10 CNN, where DCF exhibited a smaller generalization gap despite slightly lower maximum test accuracy. The smaller DCF gap therefore did not correspond to superior test performance and may partly reflect weaker fitting of the training data. The observed difference in generalization behavior consequently appears to be architecture dependent and should not be interpreted as a universal regularization property of NEL.

### 3.4 Effect of Exploration

We next examined the effect of the exploration probability *ϵ* on NEL using the MNIST MLP. As shown in Fig. 2a, performance remained relatively stable across the evaluated nonzero exploration probabilities. Mean maximum test accuracy ranged from 97.52% to 97.64% for *ϵ* ∈ [0.02, 0.20].

**Figure 2:**
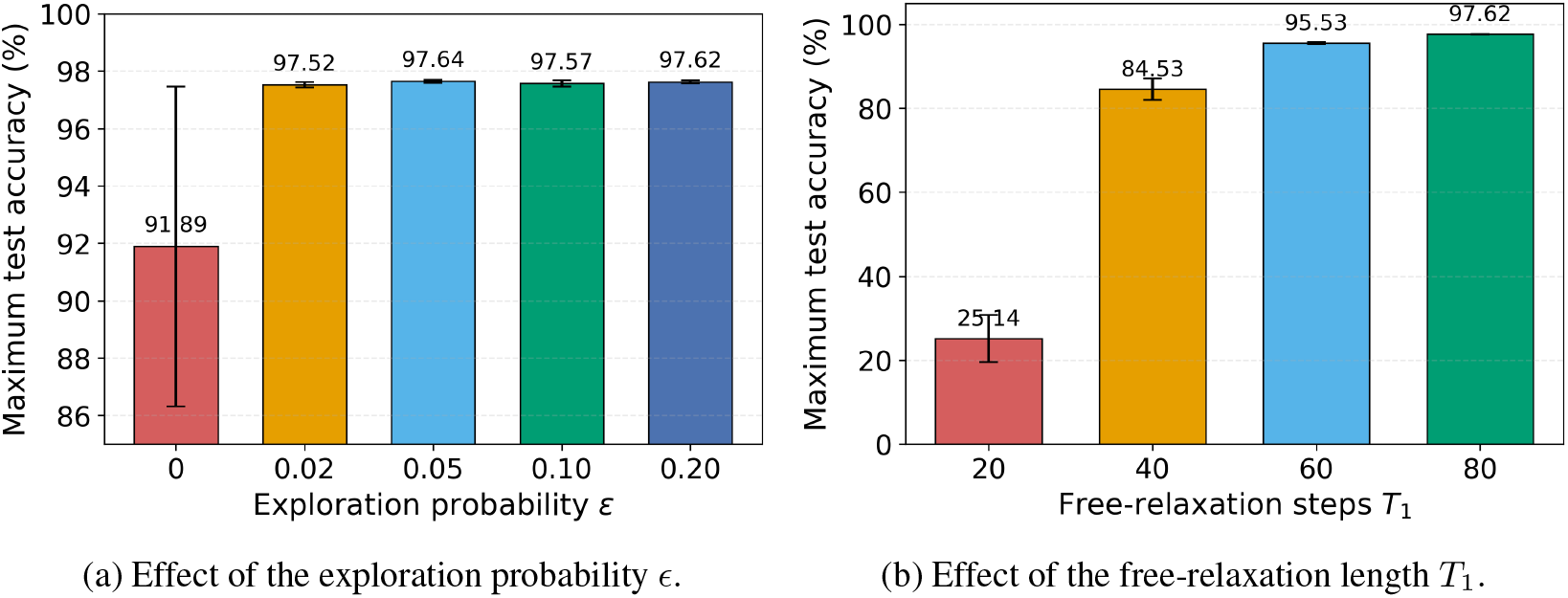
Effects of the exploration probability *ϵ* and free-relaxation length *T*_1_ on NEL performance on MNIST. Error bars indicate standard deviations across six random seeds.

In contrast, eliminating exploration substantially reduced performance. At *ϵ* = 0, accuracy decreased to 91.89 ± 5.57%, with markedly greater variability across random seeds. These results indicate that NEL is relatively insensitive to the precise exploration probability within the tested nonzero range, while some degree of exploratory action selection is important for reliable learning.

### 3.5 Effect of Free-Relaxation Length

We next examined the effect of the free-relaxation length *T*_1_. As shown in Fig. 2b, reducing *T*_1_ progressively degraded classification performance. NEL achieved 97.62 ± 0.05% at *T*_1_ = 80, compared with 95.53 ± 0.25%, 84.53 ± 2.56%, and 25.14 ± 5.60% at *T*_1_ = 60, 40, and 20, respectively. These results demonstrate that sufficient free relaxation remains important for reliable learning even though NEL eliminates the subsequent reward-nudged phase.

## 4 Discussion

We introduced Neuromodulated Equilibrium Learning (NEL), a single-equilibrium reward-modulated framework combining equilibrium relaxation (Scellier & Bengio, 2017, 2019; Ernoult et al., 2019; Laborieux et al., 2021; Laborieux & Zenke, 2022) with reward-based learning (Pozzi et al., 2020; Kubo, 2026b). Unlike conventional EP and reward-based EP (REP), NEL retains only the free equilibrium and replaces reward-nudged relaxation and contrastive learning with direct neuromodulatory feedback and local synaptic plasticity. Its distinguishing features are single equilibrium computation, sparse reward-based teaching, reciprocal feedback, and derivative-free local plasticity.

NEL remained relatively close to three-phase REP in the shallow MLPs, while larger gaps emerged for the deeper FMNIST model and CIFAR-10 CNN. Direct Cost Feedback (DCF) showed that replacing the sparse reward signal with the full supervised output error did not eliminate these differences, indicating that reward sparsity alone is insufficient to explain the performance gap. In contrast, BPTT recovered much of the performance of three-phase learning without additional nudged equilibrium phases, matching or slightly exceeding three-phase EP on the FMNIST MLPs and remaining close on MNIST, while a larger gap persisted on CIFAR-10. Although BPTT is not biologically plausible, these results suggest that performance depends more broadly on how effectively teaching information is propagated through the network dynamics rather than simply on the number of equilibrium phases.

Additional analyses further characterized NEL’s behavior. NEL showed smaller late-training train test gaps than DCF in the evaluated MLPs, but not in the CIFAR-10 CNN. Performance was robust across tested nonzero exploration probabilities, whereas eliminating exploration reduced accuracy and increased variability. Shortening the free-relaxation period also strongly degraded performance, indicating that sufficient free-equilibrium computation remains important.

Several limitations and extensions remain. Learned or approximate feedback pathways could address the weight-symmetry limitation of the current reciprocal-feedback formulation. Recurrent, heterogeneous-time-constant, and spiking architectures (Keller et al., 2024; Kubo, 2026a; Perez Nieves et al., 2021; Kubo et al., 2026; Kubo, 2026c; Martin et al., 2021) provide natural settings for richer neuronal dynamics. NEL may also support biologically plausible reinforcement learning (Kubo et al., 2022) and continual reward learning through Sleep Replay Consolidation (Kubo et al., 2025; Tadros et al., 2022). Overall, NEL demonstrates useful reward-driven learning from a single free equilibrium, while the BPTT results motivate future work on more effective biologically plausible credit propagation within a single-equilibrium framework.

## Acknowledgments

This research was enabled in part by computational resources provided by the Digital Research Alliance of Canada (alliancecan.ca).

## A Relationship Between NEL and the Nudged Dynamics of EP

The formulation of EP helps clarify the computation performed by the nudged phase and how this computation differs from the direct neuromodulatory feedback used in NEL. At equilibrium,

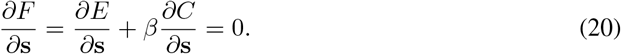

At the free equilibrium **s**^0^, *β* = 0 and therefore *∂E/∂***s** = 0. For a sufficiently small perturbation *β*, a first-order expansion around the free equilibrium gives

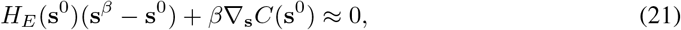

where 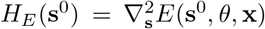 denotes the Hessian of the energy with respect to the neuronal states at the free equilibrium. Consequently,

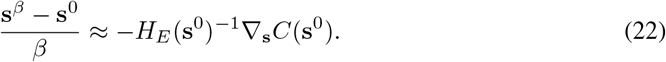

This expression illustrates that the weakly clamped phase of EP transforms the task-dependent perturbation through the local recurrent response of the network before synaptic plasticity is computed. NEL does not explicitly reproduce this relaxation-mediated transformation. Instead, it retains the reward-dependent signal at the free equilibrium and communicates it directly through reciprocal feedback connections. Specifically, the output signal 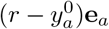 is propagated toward earlier layers and combined with local equilibrium activities to determine synaptic plasticity.

NEL should therefore not be interpreted as mathematically equivalent to the nudged dynamics of EP. Rather, it provides a simpler alternative in which reward-dependent credit is communicated directly through the feedback pathway instead of being transformed through additional nudged equilibrium relaxations. The weaker performance of NEL in deeper and convolutional architectures may therefore reflect limitations in how effectively this direct feedback pathway propagates credit through the network, although the present experiments do not isolate the responsible mechanism.

## B Experimental Details

### Activation bounds

All output layers used the hard-sigmoid activation. The MNIST and one-hidden-layer FMNIST models used ReLU hidden units. For the two-hidden-layer FMNIST models, leaky ReLU activities were bounded to [−1.2, 1.2]. For CIFAR-10, ReLU activities were bounded to [0, 1.2].

### NEL dynamics

NEL used synchronous discrete-time Euler updates during free relaxation. For the MLP implementation, let **h**_0_ = **x** denote the input, **h**_*l*_ the activity of hidden layer *l*, and **y** = **h**_*L*_ the output activity. For hidden layers *l* = 1, …, *L* − 1, the free dynamics were

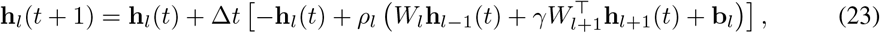

while the output dynamics were

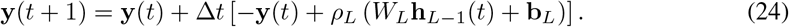

All neuronal states on the right-hand side were evaluated at time *t*, so that the updates were synchronous. The reciprocal-feedback strength was fixed at *γ* = 1.0. After free relaxation, the selected-action reward signal was propagated through the transpose feedback pathway without activation derivatives. All layer-wise synaptic updates were computed using the same free-equilibrium activities and pre-update parameters before any parameter was modified.

### Direct Cost Feedback dynamics

Direct Cost Feedback (DCF) used the same free-equilibrium relaxation, reciprocal-feedback pathway, and local synaptic update structure as NEL. The reciprocal-feedback strength was again fixed at *γ* = 1.0. Instead of the selected-action reward signal used by NEL, DCF used the full supervised output teaching signal

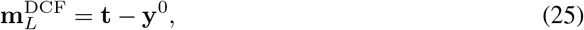

where **t** is the one-hot target vector and **y**^0^ is the output activity at the free equilibrium. This signal was propagated recursively through the transpose feedback weights without activation derivatives, using the same feedback rule and local synaptic update as NEL. DCF used no action selection, exploration, reward signal, reward-nudged phase, or contrastive update. As in NEL, all layer-wise synaptic updates were computed before any parameters were modified.

### REP dynamics

All REP experiments used the symmetric three-phase formulation. After the free phase, the network was relaxed under positive and negative reward nudging using +*β* and −*β*, respectively, and the parameter update was computed from the two nudged equilibria. The selected action and the binary reward obtained from the free equilibrium were kept fixed across both nudged phases. For CIFAR-10, the REP configuration, including *T*_1_ = 120, follows the setting used in the original REP study.

### EP dynamics

Standard Equilibrium Propagation (EP) used the same symmetric three-phase structure as REP, consisting of a free phase followed by positive and negative nudged phases using +*β* and −*β*. In contrast to REP, which nudged only the selected output according to its binary reward, standard EP used the full supervised target during both nudged phases. For the MSE experiments, the target was the one-hot class vector. The EP configurations were matched to the corresponding REP configurations in architecture, activation function, relaxation lengths, and, unless otherwise stated, optimization settings.

### BPTT dynamics

Following Laborieux et al. (2021), BPTT was applied to the same free-relaxation dynamics as the corresponding equilibrium models. The network was first evolved for *T*_1_ − *T*_2_ steps without retaining the temporal computation graph. The resulting neuronal states were then detached, and supervised gradients were propagated through the final *T*_2_ = 12 steps of the free relaxation using truncated backpropagation through time. The loss was computed from the final output activity using the full one-hot supervised target. BPTT therefore required no additional nudged equilibrium phases, while retaining gradient-based credit assignment through the final portion of the free-relaxation trajectory.

### Convolutional architecture

The CIFAR-10 model used two convolutional layers with 64 and 128 channels. Both layers used 5 × 5 kernels, stride 1, no padding, and 2 × 2 max pooling. The pooled representation was connected directly to the 10-dimensional output layer without an additional fully connected hidden layer. In both NEL and DCF, the reciprocal pathway reused the max-pooling switches for unpooling and applied transpose convolution with tied forward kernels.

### Optimization and reporting

REP, EP, and BPTT used SGD with zero momentum. For the CIFAR-10 REP, EP, and BPTT experiments, weight decay of 3 × 10^−4^ was applied to each trainable layer. No learning-rate schedule was used in the reported experiments. All reported results were averaged over six random seeds. BPTT learning rates were tuned separately for each architecture. Model-specific learning rates, relaxation lengths, and training durations are summarized in Table 2. Complete implementation details and experiment configurations are provided in the accompanying code repository.

**Table 2:** Principal model and training configurations for NEL, Direct Cost Feedback (DCF), BPTT, three-phase REP, and three-phase EP. *T*_1_ denotes the free-relaxation length. For REP and EP, *T*_2_ denotes the length of each nudged relaxation; for BPTT, *T*_2_ denotes the number of final free-relaxation steps through which gradients were backpropagated.

| Dataset | Method | Architecture | Activation | $T_1$ | $T_2$ | $\Delta t$ | LRs | Ep. |
| --- | --- | --- | --- | --- | --- | --- | --- | --- |
| MNIST | NEL | 784-512-10 | ReLU | 80 | – | 0.1 | 0.1/0.1 | 80 |
| MNIST | DCF | 784-512-10 | ReLU | 80 | – | 0.1 | 0.1/0.1 | 80 |
| MNIST | BPTT | 784-512-10 | ReLU | 80 | 12 | – | 0.4/0.04 | 80 |
| MNIST | REP (3-phase) | 784-512-10 | ReLU | 80 | 12 | – | 0.3/0.02 | 80 |
| MNIST | EP (3-phase) | 784-512-10 | ReLU | 80 | 12 | – | 0.3/0.02 | 80 |
| FMNIST | NEL | 784-1024-10 | ReLU | 80 | – | 0.1 | 0.1/0.1 | 80 |
| FMNIST | DCF | 784-1024-10 | ReLU | 80 | – | 0.1 | 0.1/0.1 | 80 |
| FMNIST | BPTT | 784-1024-10 | ReLU | 80 | 12 | – | 0.21/0.0011 | 80 |
| FMNIST | REP (3-phase) | 784-1024-10 | ReLU | 80 | 12 | – | 0.2/0.01 | 80 |
| FMNIST | EP (3-phase) | 784-1024-10 | ReLU | 80 | 12 | – | 0.2/0.01 | 80 |
| FMNIST | NEL | 784-512-256-10 | Bounded<br>LeakyReLU | 100 | – | 0.1 | $0.12 \times 3$ | 120 |
| FMNIST | DCF | 784-512-256-10 | Bounded<br>LeakyReLU | 100 | – | 0.1 | $0.08 \times 3$ | 120 |
| FMNIST | BPTT | 784-512-256-10 | Bounded<br>LeakyReLU | 100 | 12 | – | $0.1 \times 3$ | 120 |
| FMNIST | REP (3-phase) | 784-512-256-10 | Bounded<br>LeakyReLU | 100 | 12 | – | 0.2/0.08/0.02 | 120 |
| FMNIST | EP (3-phase) | 784-512-256-10 | Bounded<br>LeakyReLU | 100 | 12 | – | 0.2/0.08/0.02 | 120 |
| CIFAR-10 | NEL | Conv64-Conv128-10 | Bounded ReLU | 80 | – | 0.2 | $0.02 \times 3$ | 120 |
| CIFAR-10 | DCF | Conv64-Conv128-10 | Bounded ReLU | 80 | – | 0.2 | $0.02 \times 3$ | 120 |
| CIFAR-10 | BPTT | Conv64-Conv128-10 | Bounded ReLU | 120 | 12 | – | 0.13/0.05/0.001 | 120 |
| CIFAR-10 | REP (3-phase) | Conv64-Conv128-10 | Bounded ReLU | 120 | 12 | – | 0.13/0.05/0.001 | 120 |
| CIFAR-10 | EP (3-phase) | Conv64-Conv128-10 | Bounded ReLU | 120 | 12 | – | 0.13/0.05/0.001 | 120 |

For the two-hidden-layer FMNIST DCF experiment, using the NEL-matched learning rate of 0.12 for each layer resulted in late-training instability and eventual collapse. The reported stable configuration therefore used a learning rate of 0.08 for each layer. BPTT learning rates were tuned separately for each architecture.

## Notes

### Competing Interest Statement

The authors have declared no competing interest.

### Summary of Updates

Added standard three-phase Equilibrium Propagation (EP) and truncated BPTT baselines, and revised the analysis and discussion accordingly.

